# Analysis of Nutritional Components in Brown Sugar and Its Effects on Anti-fatigue, Alleviation of Qi Deficiency and Blood Stasis, and Antioxidant Effects

**DOI:** 10.1101/2025.09.15.676350

**Authors:** Zhang Qiang, Yue Shuai, Zhang Sicong, Li Yi, Xu Guanghui, Wan Qiang, Wang Bao, Qiu Shiming, Zhang Bowei, Huang Yinru, Wang Jian

**Affiliations:** Guangxi Key Laboratory for High-value Utilization of Manganese Resources, School of Chemistry and Materials Science, Guangxi Minzu Normal University, Chongzuo, Guangxi, China; COFCO Chongzuo Sugar Industry Co., Ltd./Guangxi Sugarcane Industry Chain Extension Engineering Research Center, Chongzuo, Guangxi, China; COFCO Nutrition and Health Research Institute Co, Ltd./Beijing Engineering Laboratory of Geriatric Nutrition & Foods/Beijing Key Laboratory of Nutrition & Health and Food Safety, Beijing, China; COFCO Sugar Co., Ltd./Key Laboratory of Quality & Safety Control for Sugar Crops and Tomato, Ministry of Agriculture and Rural Affiairs/National Sugar Processing TechnologyResearch and Development Center, Changji, Xinjiang, China

**Keywords:** Brown sugar, nutritional components, anti fatigue, qi deficiency and blood stasis, antioxidant

## Abstract

Brown sugar (BS), Red granulated sugar (RGS) and White sugar(WS) are three key sweeteners and food additives in China. Notably, BS possesses significant nutritional value and has been widely used in Chinese medicine for its supportive therapeutic properties.This study evaluated the anti-fatigue, the amelioration of qi deficiency and blood stasis, as well as the antioxidant activities of these sugars using a zebrafish model. BS and RGS demonstrated significantly greater efficacy than WS in terms of both anti-fatigue effects and amelioration of qi Deficiency and blood stasis in the zebrafish model. Antioxidant assays revealed that BS, RGS, and WS (500 μg/mL) significantly enhanced superoxide dismutase (SOD) activity in zebrafish embryos compared with the control group. Furthermore, the mineral composition of sugar were determined. Nine mineral elements—potassium (K), calcium (Ca), sodium (Na), magnesium (Mg), iron (Fe), zinc (Zn), copper (Cu), phosphorus (P), and selenium (Se)—were analyzed. BS exhibited markedly higher concentrations of P, Ca, Fe, and Mg than RGS and WS, while selenium (detected at 50 mg/kg) was exclusively detected in BS. Intriguingly, RGS contained four-fold higher potassium levels than BS. Total polyphenol content, quantified by Folin-Ciocalteu assay, was 4.35 g/kg in BS and 1.53 g/kg in RGS, with no detectable polyphenols in WS.

## Introduction

Sugar serves as a critical food ingredient and a primary energy source for the human body. Currently, the predominant sugarcane-derived sugars used as sweeteners in China include brown sugar(BS), red granulated sugar (RGS), and white sugar(WS) [1, 2]. Among these, BS, recognized as both a traditional Chinese food and significant sweetener additive, possesses a long history of widespread use.It is valued not only for its distinctive sweet flavor, but also for its rich nutritional composition and potential health benefits [3,4,5,6]. Traditional Chinese Medicine (TCM) records attribute effects such as moistening lung qi, tonifying the five zang organs, promoting fluid production, detoxification, fortifying the spleen, and soothing liver qi to brown sugar.. In China, BS is commonly utilized as an energy supplement for physically debilitated individuals,, as well as for women during menstruation and lactation. However, existing research on BS has predominantly focused on physicochemical characterization. Information regarding the biological activities of sugars and their potential correlations with nutritional components (e.g., minerals, vitamins, organic acids, amino acids, phenolic substances) remains scarce.

Zebrafish, an emerging biological model organism, offers unique advantages that facilitate its widespread application in bioactivity evaluation [7]. The zebrafish genome exhibits highly similarity to human genome, and many disease models can be effectively simulated in zebrafish, providing a reliable platform for drug screening and mechanistic studies [8].

This study pioneers the use of a zebrafish model to evaluate BS for its anti-fatigue effects, its capacity to ameliorate qi deficiency and blood stasis, and its antioxidant properties. Furthermore, correlations between the nutritional constituents of BS and its observed bioactivities were investigated. This work establishes a feasible methodology for assessing the functional efficacy of sugars and elucidates potential connections between the bioactivity of BS and its nutritional profile. The results could contribute to the enhanced utilization of sugar and provide fundamental data for the development and application of functional foods based on BS.

## 2. Material and methods

### 2.1. Materials and devices

#### 2.1.1 Materials and Regents

BS was produced by COFCO Beihai Sugar Industry Co., Ltd. RGS was produced by Yuebei Sugar Industry Co., Ltd. WS was produced by COFCO Chongzuo Sugar Industry Co., Ltd.

Zebrafish were maintained in system water at 28°C. The system water formulation consisted of 200 mg of instant sea salt added per liter of reverse osmosis water, yielding water with the following parameters: conductivity 450-550 μS/cm; pH 6.5-8.5; hardness 50-100 mg/L (as CaCO_3_ equivalent). System water and zebrafish were provided by Huante Biotechnology Co., Ltd. The Experimental Animal Use License Number is SYXK (Zhejiang) 2022-0004. Husbandry practices complied with the requirements of AAALAC International accreditation (Certification number: 001458). The Institutional Animal Care and Use Committee (IACUC) ethics review number is IACUC-2024-f9114-01.

Salidroside was sourced from Huazhong Haiwei (Beijing) Gene Technology Co., Ltd. Naoxintong capsules were obtained from Shanxi Buchang Pharmaceutical Co., Ltd.. N-acetyl-L-cysteine (NAC) was obtained from Shanghai Aladdin Biochemical Technology Co., Ltd. N-Acetyl-L-cysteine (NAC), hydrogen peroxide, nitric acid (superior purity grade), anhydrous sodium sulfite, methylcellulose, menadione, and dimethyl sulfoxide were procured from Shanghai Aladdin Biochemical Technology Co., Ltd., China. Isoprenaline hydrochloride (KIWNH-SG) was sourced from TCI, China. Single-element standard solutions (1000 μg/mL) for Na, Mg, P, K, Ca, Mn, Fe, Cu, Zn, and Se were obtained from the National Nonferrous Metals and Electronic Materials Analysis and Testing Center, China. Ultra-pure water (resistivity 18.2 MΩ · cm) was utilized. CellROX™ Green Reagent was purchased from Invitrogen, USA. The BCA protein concentration assay kit was sourced from Bosch Bioengineering Co., Ltd., China. The Total Superoxide Dismutase (SOD) Assay Kit (WST-1 Method; Product No. A001-3-2) was obtained from the Nanjing Institute of Bioengineering, China. Sodium chloride injection (0.9%) was obtained from Hunan Kelun Pharmaceutical Co., Ltd., China.

#### 2.1.2 Instruments and devices

JY ULTIMA 2 Inductively Coupled Plasma Optical Emission Spectrometer (ICP-OES; Horiba Jobin Yvon S.A.S., France). Mars 6 Classic Microwave Digestion System (CEM Corporation, USA). Milli-Q Ultrapure Water System (Merck Millipore, USA). AX224ZH/E Analytical Balance (Ohaus International Trade (Shanghai) Co., Ltd., China). SZX7 Stereo Microscope (Olympus, Japan). VertA1 Charge-Coupled Device (CCD) Camera (Shanghai Tusen Vision Technology Co., Ltd., China). CP214 Precision Electronic Balance (OHAUS, USA). 6-Well Cell Culture Plates (Zhejiang Belambo Biotechnology Co., Ltd., China). ZebraLab 3.22.3.31 Behavior Tracking System (Viewpoint, France). 96-Well Plates (Nest Biotech, China). JXFSTPRP-24L Automatic Sample Homogenizer/Grinder (Shanghai Jingxin Industrial Development Co., Ltd., China). PIC017/21 High-Speed Refrigerated Centrifuge (Thermo Fisher Scientific, USA). AZ100 Motorized Focusing Continuous Zoom Fluorescence Microscope (Nikon, Japan). SPARK Multimode Microplate Reader (Tecan Group Ltd., Austria)

### 2.2 Relieves physical fatigue effect

#### 2.2.1 Determination of MTC

Wild-type AB strain of zebrafish with 4 days of post-fertilization (4 dpf) were randomly selected and distributed into 6-well plates (30 zebrafish per well in 3 mL of water). As listed in Table 1, experimental groups included: control, model, BS, RGS, and WS. After 24 h incubation at 28°C, a physical fatigue model was induced in all groups except the control by exposure to anhydrous sodium sulfite. The Maximum Tolerated Concentration (MTC) of test samples on model zebrafish was determined following 30 min treatment at 28°C.

**Table 1.**
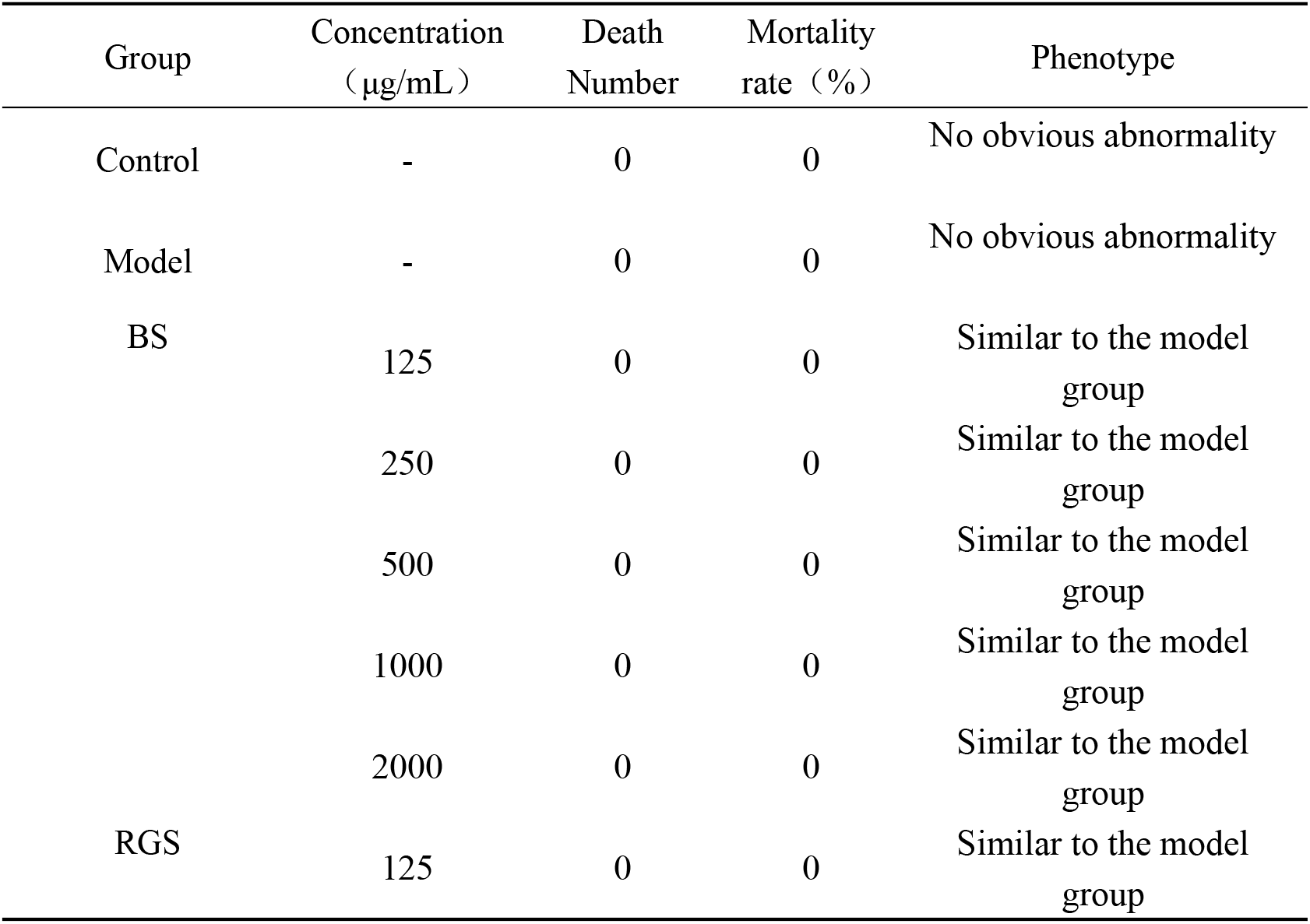
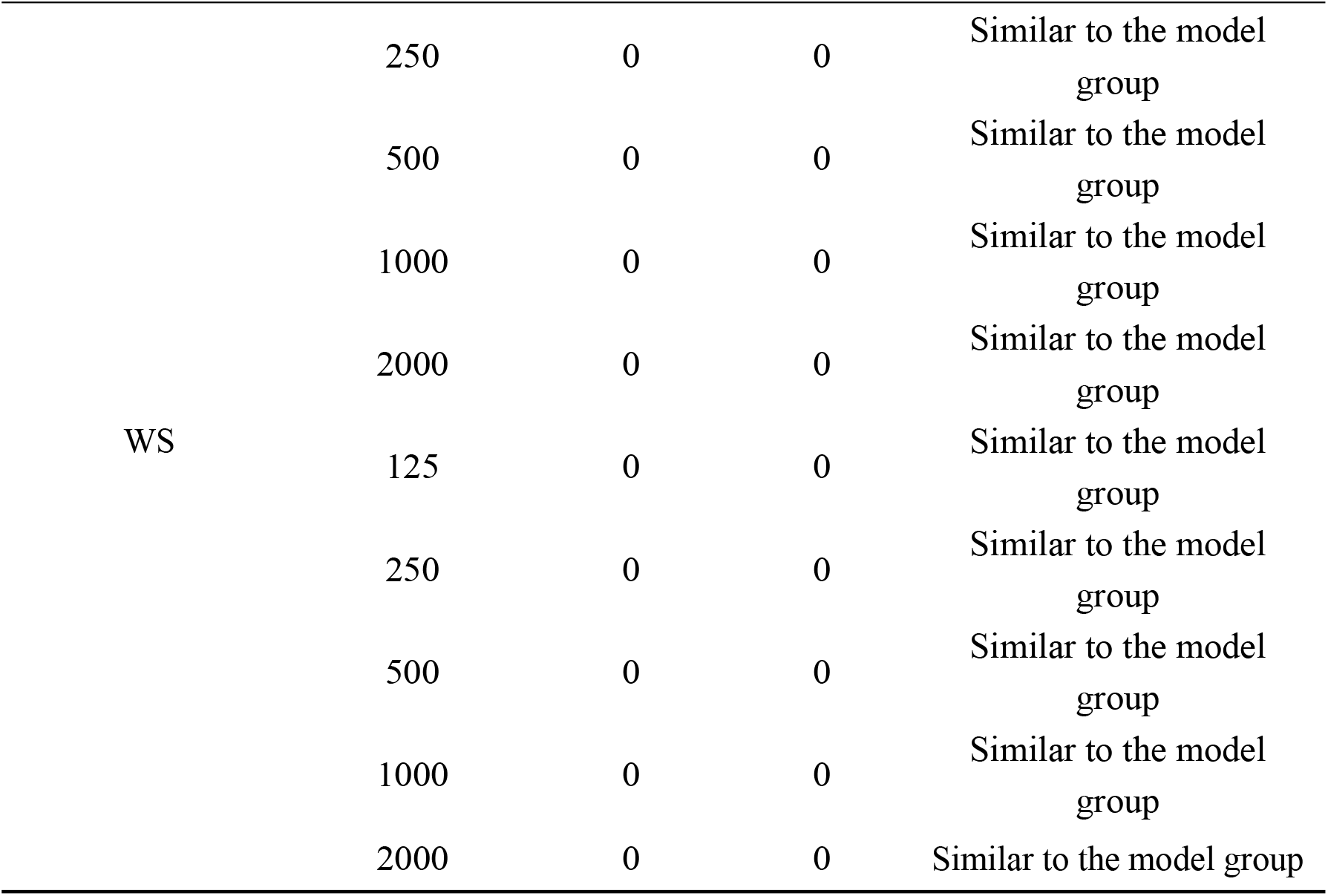
MTC for relieving physical fatigue of samples(n = 30)

#### 2.2.2 Relieving physical fatigue effect

Wild-type AB strain zebrafish (4 dpf) were randomly allocated to 6-well plates (30 zebrafish per well in 3 mL of water). As specified in Table 1, groups included: control, model, positive control (4000 μg/mL salidroside), BS, RGS, and WS. After 24 h at 28°C, a physical fatigue model was induced in all groups except the control via anhydrous sodium sulfite exposure. Subsequently, 10 zebrafish per experimental group were individually transferred to a 96-well plate (1 zebrafish/well, 200 μL/well). Locomotor activity was recorded for 30 min using a behavior analyzer, with total movement distance quantified. Statistical analysis of this endpoint evaluated the efficacy of samples in alleviating physical fatigue.

### 2.3 Relieves qi deficiency and blood stasis effect

#### 2.3.1 Determination of MTC

The MTC determination followed the protocol outlined in Section 2.2.1, with sample details provided in Table 2. Immediately after group allocation, a qi deficiency and blood stasis model (per traditional Chinese medicine theory) was induced in all groups except the control by co-exposure to anhydrous sodium sulfite and isoprenaline hydrochloride. The MTC was determined after 48 h treatment at 28°C.

**Table 2.**
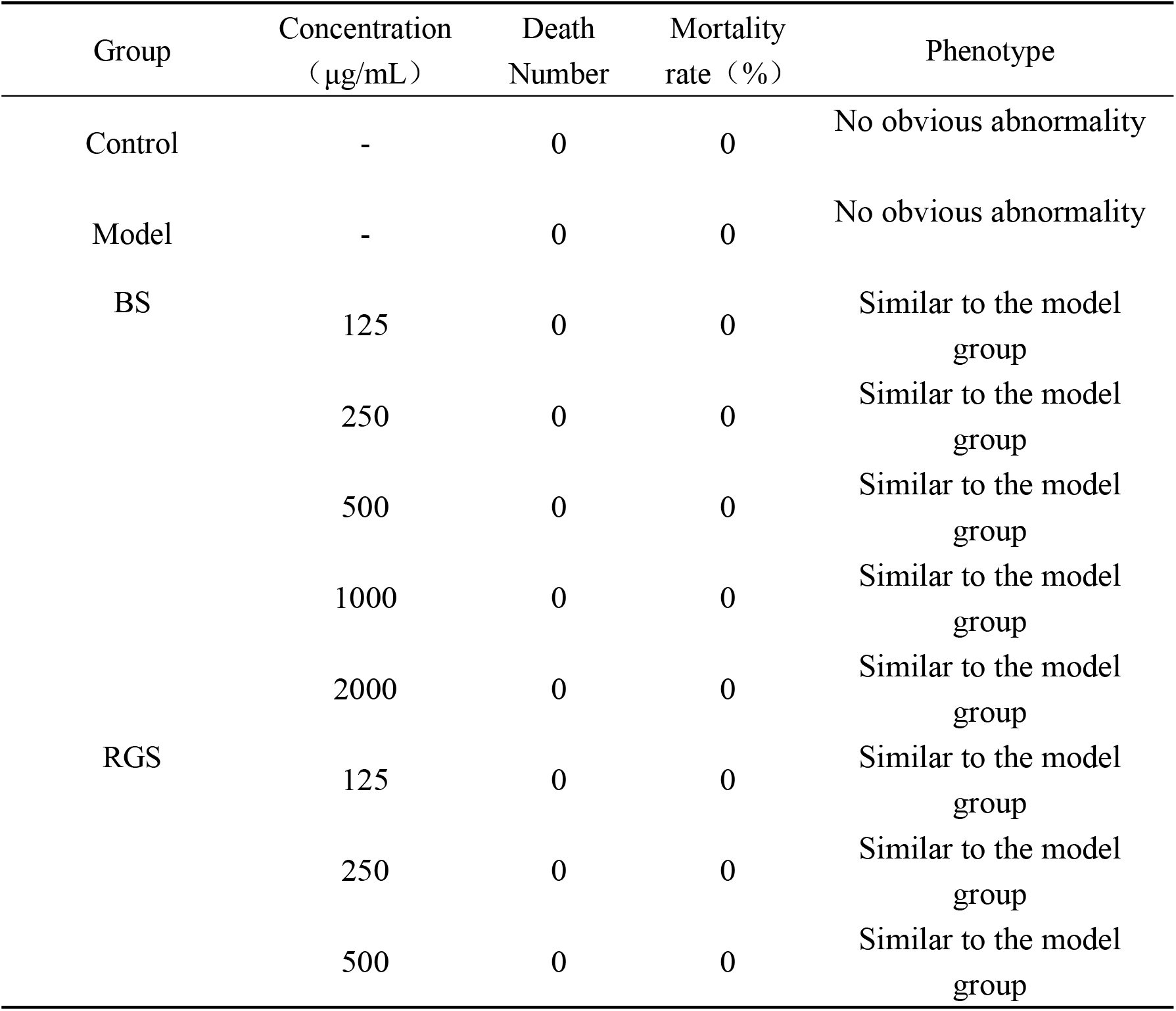
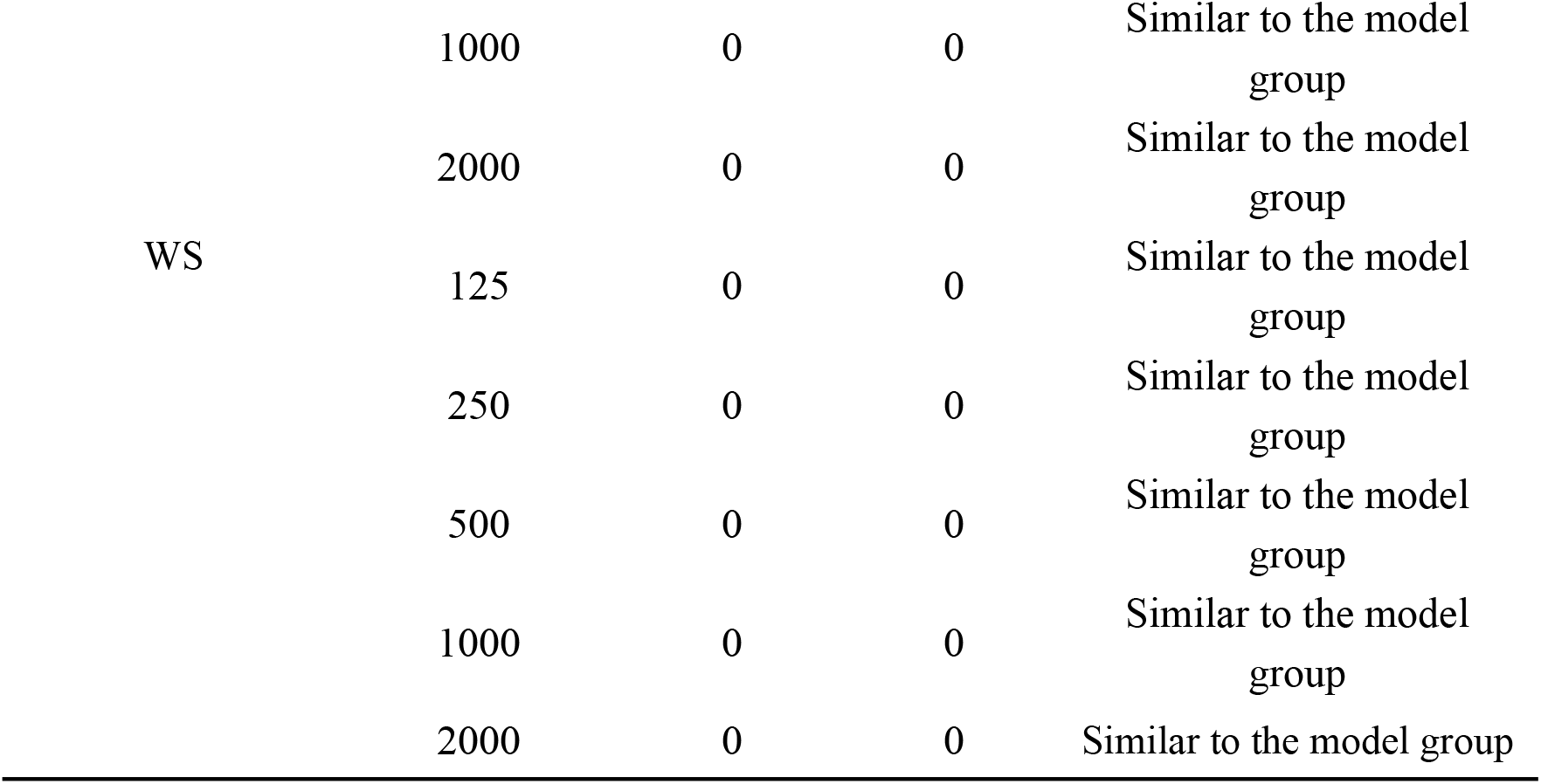
MTC for improving qi deficiency and blood stasis of samples(n = 30)

#### 2.3.2 Relieving qi deficiency and blood stasis effect

Groups included: water-soluble test samples, positive control (31.2 μg/mL Naoxintong capsules), control, and model (3 mL water/well). A qi deficiency and blood stasis model was induced in all groups except the control via co-exposure to anhydrous sodium sulfite and isoprenaline hydrochloride. After 48 h at 28°C, 10 zebrafish per group underwent cardiac function assessment using a heartbeat blood flow analysis system. Blood flow velocity and cardiac output were measured, with statistical analysis of these parameters evaluating efficacy against qi deficiency and blood stasis.

### 2.4 Antioxidant effect (Determination of SOD)

#### 2.4.1 Determination of MTC

MTC determination replicated Section 2.2.1 methodology (sample details in Table 3). After 2 h pre-treatment at 28°C, an oxidative damage model was induced in all groups except the control via menadione exposure. The MTC was determined after 22 h incubation at 28°C.

**Table 3.**
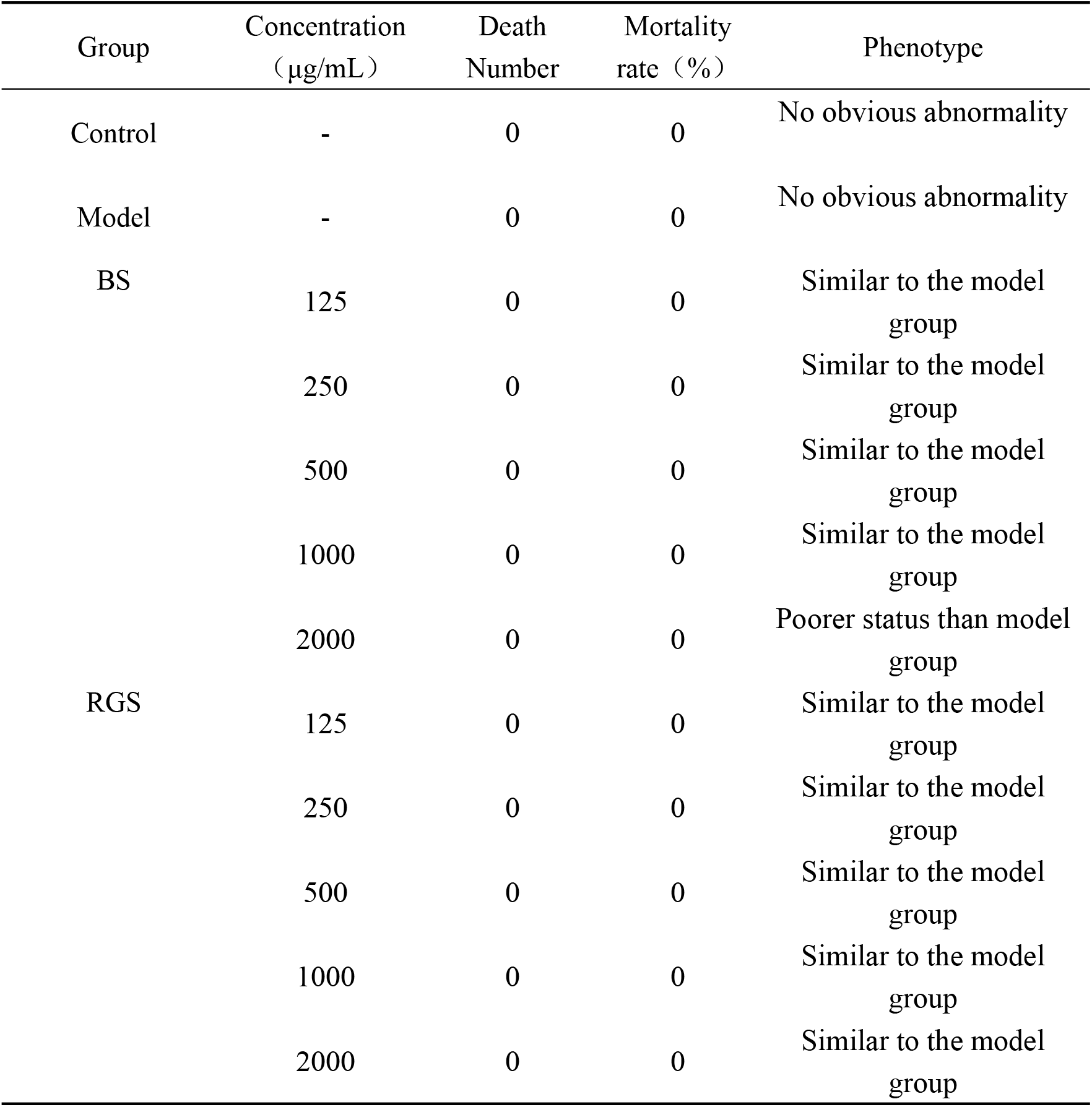
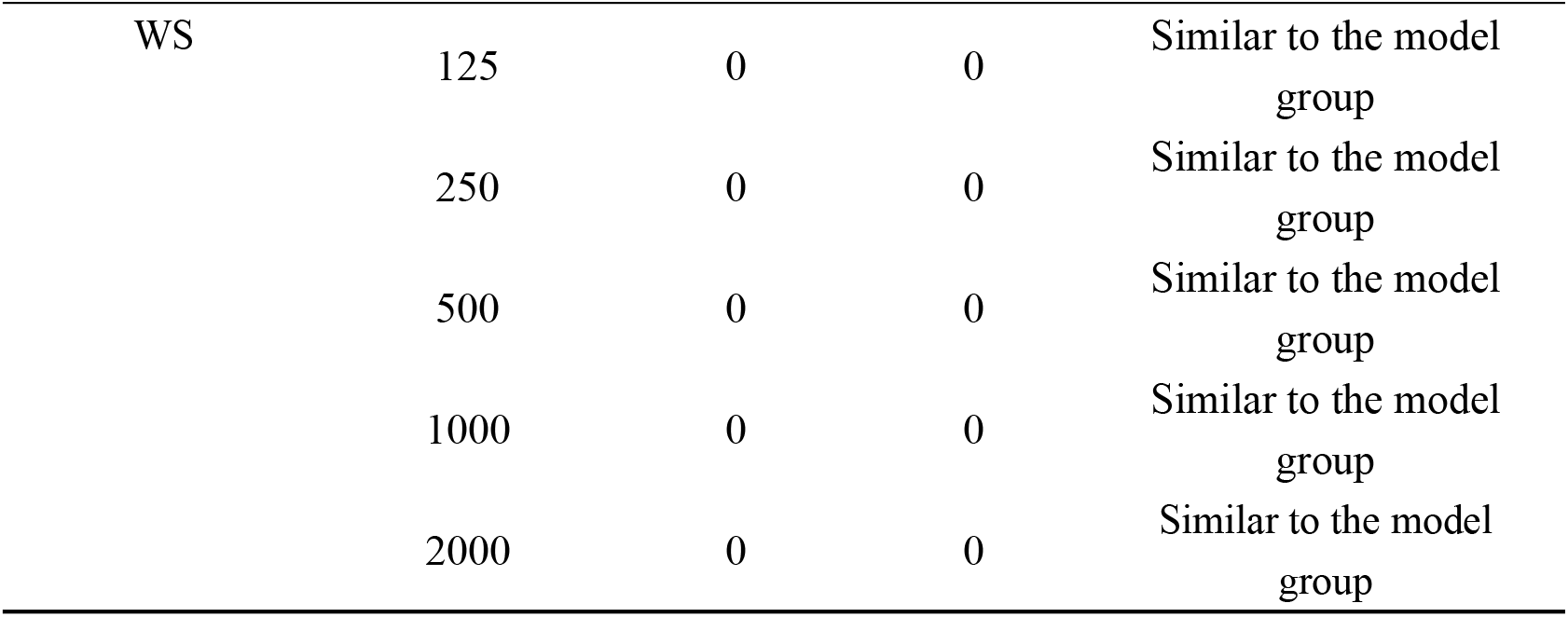
MTC for antioxidant effect of samples(n = 30)

#### 2.4.2 Antioxidant effect(Yolk sac fluorescence intensity)

Albino strain zebrafish larvae (melanin allele mutant; 3 dpf) were randomly assigned to 6-well plates (30 larvae/well). Groups included: water-soluble samples, positive control (62.5 μg/mL NAC), control, and model (3 mL water/well). After 2 h at 28°C, oxidative damage was induced via menadione exposure (all groups except control). Following 22 h at 28°C, larvae were stained with CellROX® for 30 min and washed 3 times with standard dilution water. Ten larvae per group were imaged using fluorescence microscopy. Antioxidant efficacy was evaluated by statistical analysis of yolk sac fluorescence intensity.

#### 2.4.3 Antioxidant effect (SOD vitality)

Wild-type AB strain zebrafish larvae (3 dpf) were randomly assigned to 6-well plates (30 larvae/well in 3 mL water; n=3 replicates). Groups included water-soluble samples (Table 3), positive control (62.5 μg/mL NAC), control, and model. After 2 h at 28°C, oxidative damage was induced via menadione exposure (all groups except control). Post 22 h incubation (28°C), larvae were homogenized. Protein concentration was determined using a BCA assay kit. SOD activity was quantified using an SOD detection kit and a multimode microplate reader according to manufacturer protocols. Statistical analysis of SOD activity evaluated antioxidant efficacy. Data are expressed as mean ± SE. SPSS 26.0 software performed statistical analyses; *p* < 0.05 indicated statistical significance.

### 2.5 Determination of Inorganic nutrient elements and Total Polyphenols Content

#### 2.5.1 Measurement of Inorganic nutrient elements

##### Sample Preparation

Sugar samples (0.5 g) were digested in polytetrafluoroethylene vessels with 5 mL HNO_3_ and 1 mL H_2_O_2_ (capped, 24 h room temperature). Samples underwent 10 min pre-digestion at 100°C in an acid removal system, followed by microwave digestion: 120°C (10 min), 150°C (5 min), 180°C (5 min). Digestates were evaporated to ∼1 mL at 180°C in the acid removal system, diluted to 25 mL with ultrapure water, and stored 24 h prior to analysis.

##### Standard Solutions

Mixed Standard Solution #1 (K, Ca, Na, Mg, Fe, Zn, Cu, P, Se) was serially diluted with 1% HNO_3_ to concentrations of 0, 1.00, 2.00, 5.00, 10.00, 20.00, and 50.00 µg/mL. Mixed Standard Solution #2 (Mn, Fe, Cu, Zn, Se) was diluted to 0, 0.15, 0.25, 0.30, 0.35, 0.50, 0.80, 1.00, and 2.00 µg/mL. ICP-OES operating parameters: RF power 1000 W, cooling gas 14.0 L/min, carrier gas 1.30 L/min, auxiliary gas 0.2 L/min, pump speed 20 rpm. Elemental standard curves were generated, and sample concentrations calculated accordingly.

#### 2.5.2 Polyphenol standard curve

Gallic acid standard (accurately weighed) was dissolved in distilled water to 1.5 mg/mL. Following GB/T 8313-2018 [9], aliquots (0, 0.15, 0.23, 0.35, 0.45, 0.55, 0.66 mL) were transferred to 10 mL volumetric flasks, diluted to 3 mL, mixed with 1.5 mL Folin-Ciocalteu reagent and 2 mL 5% Na_2_CO_3_, brought to volume, and incubated 20 min in darkness. Absorbance was measured at 747 nm. The calibration curve (y = 0.00574x + 0.0021; R^2^ = 0.9991, n=7) related gallic acid concentration (x, μg/mL) to absorbance (y).

#### 2.5.3 Measurement of samples

Sugar sample solution (1 mL) was transferred to a 10 mL brown volumetric flask (triplicate preparations). Absorbance was measured as above. Total polyphenol content was calculated using Equation 1:

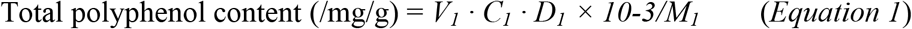

Where: *V*_*1*_ = volume of test solution (mL), *C*_*1*_ = polyphenol concentration from standard curve (μg/mL), *D*_*1*_ = dilution factor, *M*_*1*_ = sample mass (g).

### 2.6 Data analysis

Data represent mean ± standard error (SE) of three independent replicates. Statistical analyses and graphing used SPSS 26.0 and Origin 9 Pro, respectively. Significance was defined as *p* < 0.05.

## 3. Results and discussion

### 3.1 Determination of MTC

Minimum Toxic Concentration (MTC) is defined as the lowest concentration of a drug at which toxic reactions are observed in drug safety evaluations. Variability in MTC across different experimental models is a critical consideration for rational drug selection and the design of treatment protocols.

As presented in Table 1, the MTC values of BS, RGS, and WS for alleviating physical fatigue in a zebrafish anti-fatigue model were 2000 μg/mL. Similarly, as shown in Table 2, the MTC values of BS, RGS, and WS for mitigating qi deficiency and blood stasis in the same zebrafish model were also 2000 μg/mL. According to Table 3, the MTC values for the antioxidant effects of BS, RGS, and WS were 2000 μg/mL in the zebrafish anti-fatigue model.

Under the experimental conditions applied in this study, the MTC for the antioxidant effect was 2000 μg/mL for both RGS and WS, whereas that of BS was 1000 μg/mL. Consequently, the maximum dosage for all three samples was set at 2000 μg/mL in the anti-fatigue and qi deficiency and blood stasis activity assays. In contrast, for the antioxidant activity assay, the maximum dosage was 2000 μg/mL for RGS and WS, and 1000 μg/mL for BS.

### 3.2 Anti-fatigue effect of sugar

The total movement distance is a well-established metric for evaluating anti-fatigue effects in zebrafish models [10, 11]. As shown in Fig. 1 and 2, zebrafish in the model control group exhibited a significant reduction in locomotion compared to those in the normal control group, confirming the successful establishment of the fatigue model.

**Fig. 1.**
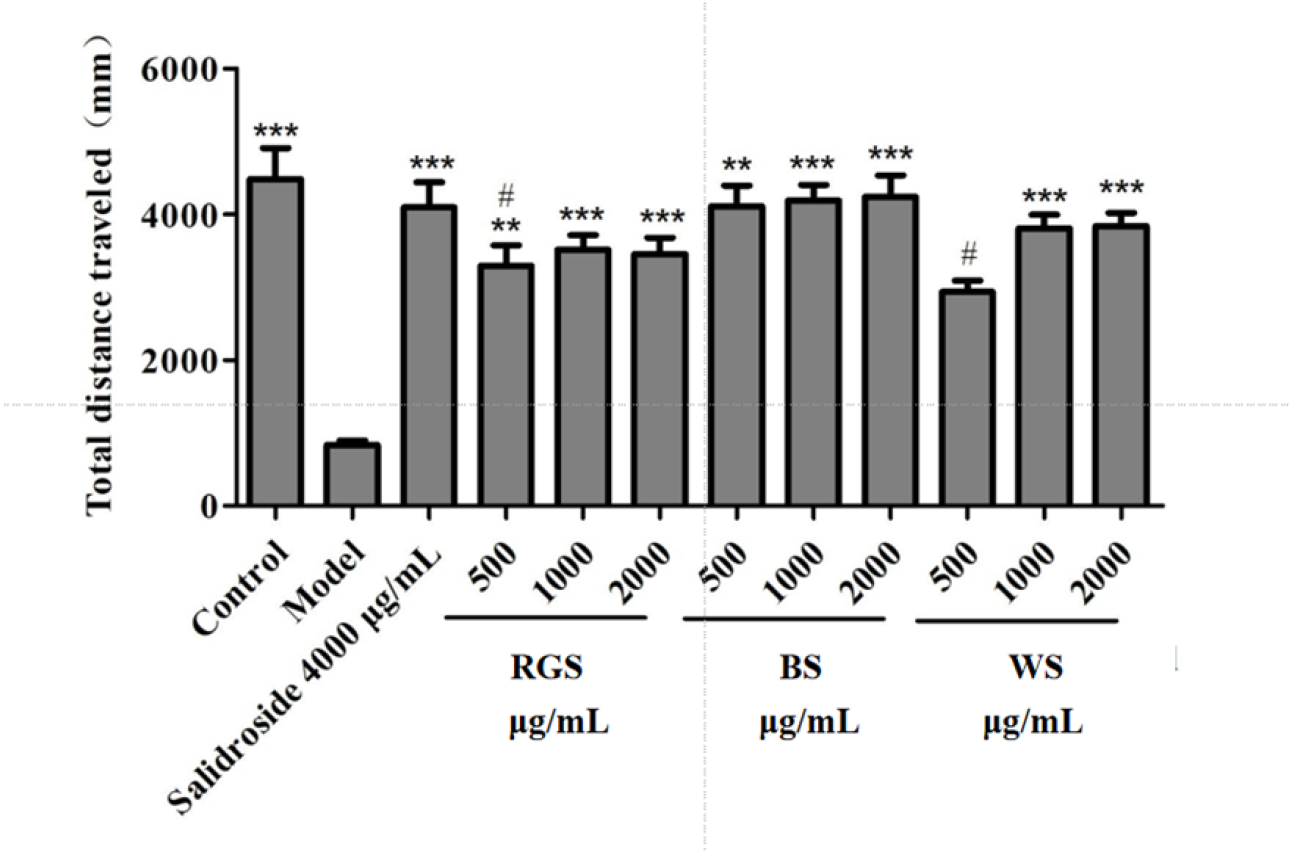
Total movement distance of zebrafish after sample processing Compared with the model control group, **p < 0.01, ***p < 0.001 Compared with the 500 µg/mL group of inganose, #p < 0.05

**Fig. 2.**
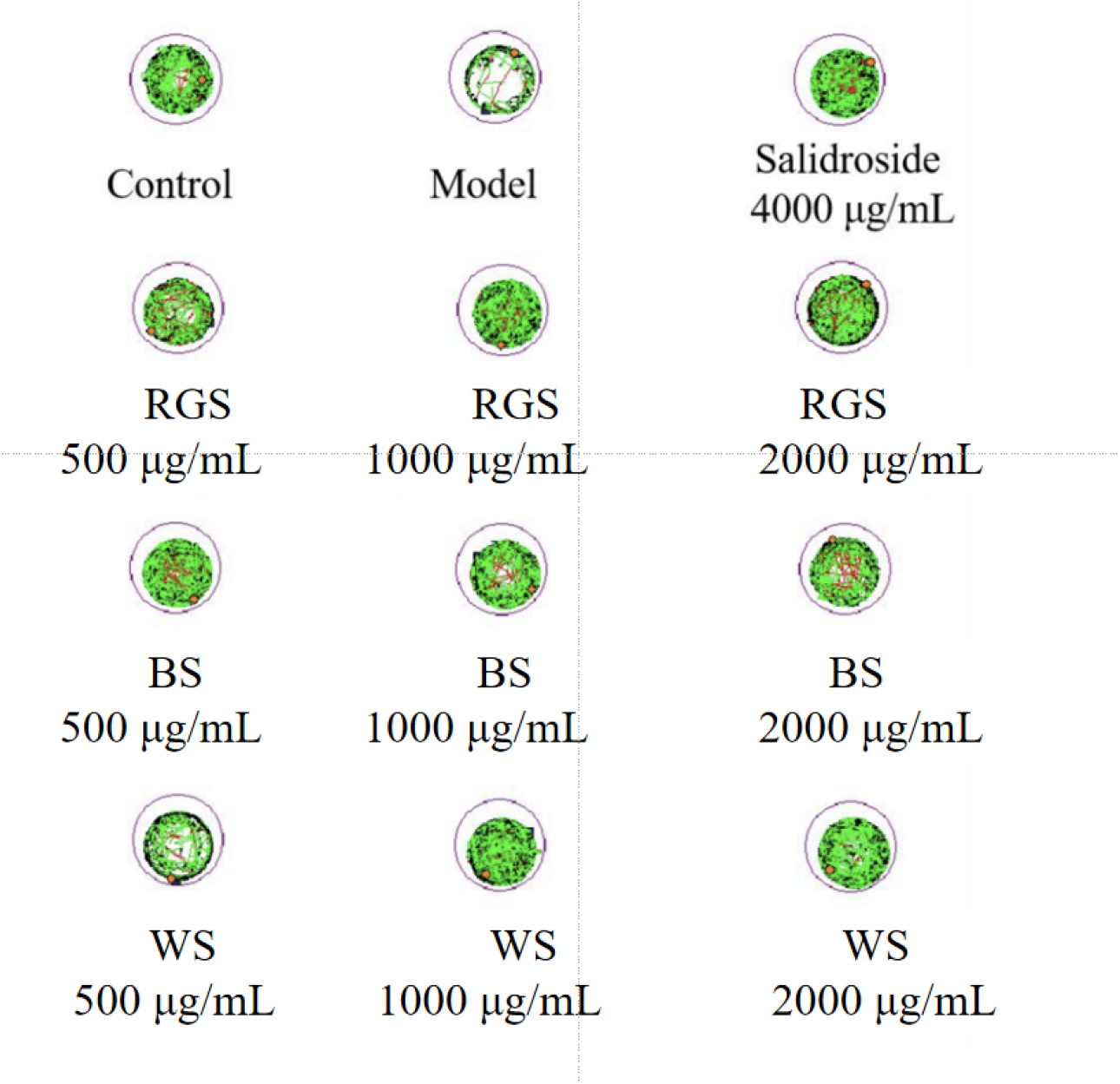
Typical diagram of zebrafish behavior trajectory after sample treatment The black line is the slow motion distance, the green line is the medium -speed movement distance, the red line is the fast movement distance

Zebrafish treated with each of the three types of sugar showed a significantly greater total movement distance than those in the model group. These results indicate that BS, RGS, and WS all exerted anti-fatigue effects under the experimental conditions. Moreover, at equivalent doses, the exercise distance of zebrafish treated with BS was longer than that with RGS or WS, suggesting a superior anti-fatigue effect of BS. At the concentration of 2000 μg/mL, the movement distance of BS-treated zebrafish was the longest among all groups, exceeding that of the salidroside positive control group by 3% at the same dose.

The anti-fatigue effects observed for BS, RGS, and WS may be attributed to their high content of sucrose and glucose, which can be directly absorbed by pancreatic beta cells in zebrafish and converted into ATP, facilitating rapid energy replenishment. Following exercise or fatigue, sugars promote muscle glycogen synthesis, reduce muscle protein breakdown, and accelerate physical recovery [12]. Monosaccharides such as glucose can directly supply energy for muscle activity, thereby alleviating fatigue associated with energy deficiency [13]. BS and RGS exhibited stronger anti-fatigue effects than WS, possibly due to the retention of more trace elements resulting from their less refined processing.

### 3.3 Relieves qi deficiency and blood stasis effect

As shown in Figs. 3 and 4, zebrafish in the model control group exhibited significantly reduced blood flow velocity and cardiac output compared to the normal control group (P < 0.05), confirming the successful establishment of the qi deficiency and blood stasis model. Relative to the model control group, both blood flow velocity and cardiac output were significantly increased in the Naoxintong capsule control group (P < 0.05). All sugar intervention groups (BS, RGS, WS) also showed markedly elevated blood flow velocity and cardiac output (P < 0.05).

**Fig. 3.**
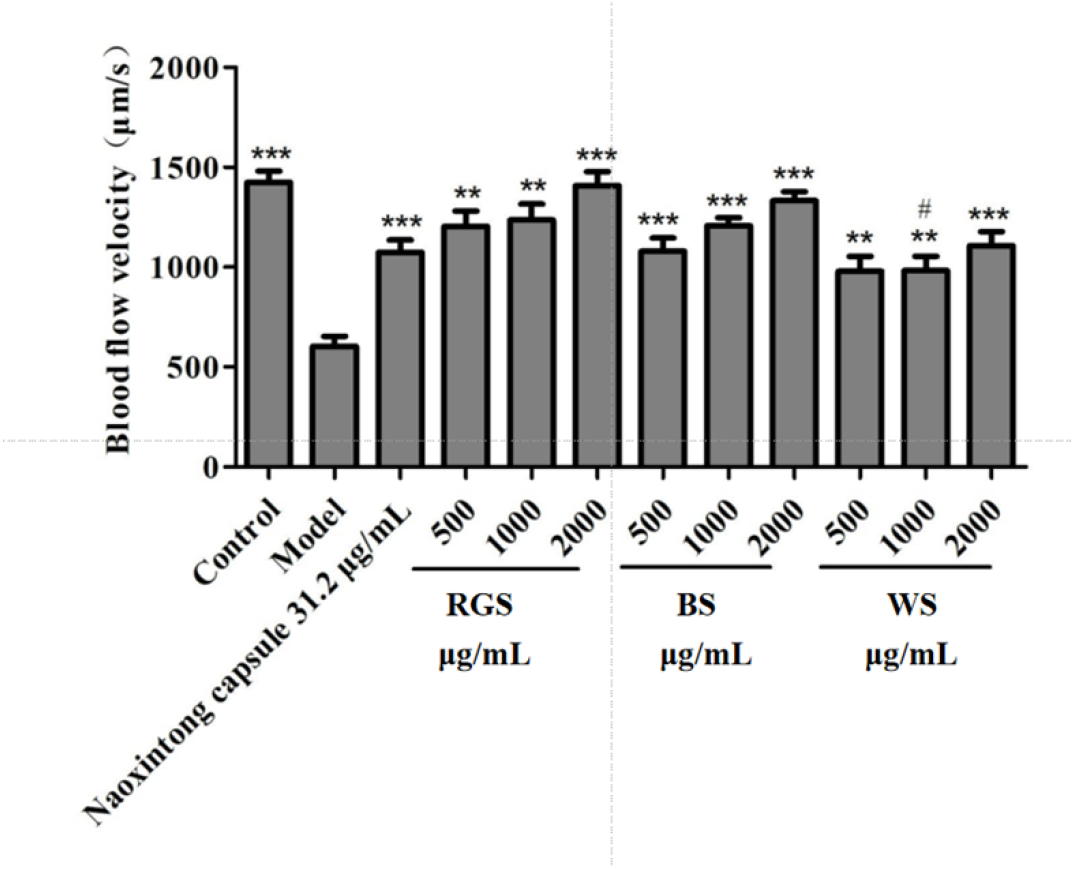
Blood flow velocity of zebrafish after sample treatment Compared with the model control group, **p < 0.01, ***p < 0.001 Compared with the 1000 µg/mL concentration group of ingamisin, #p < 0.05

**Fig. 4.**
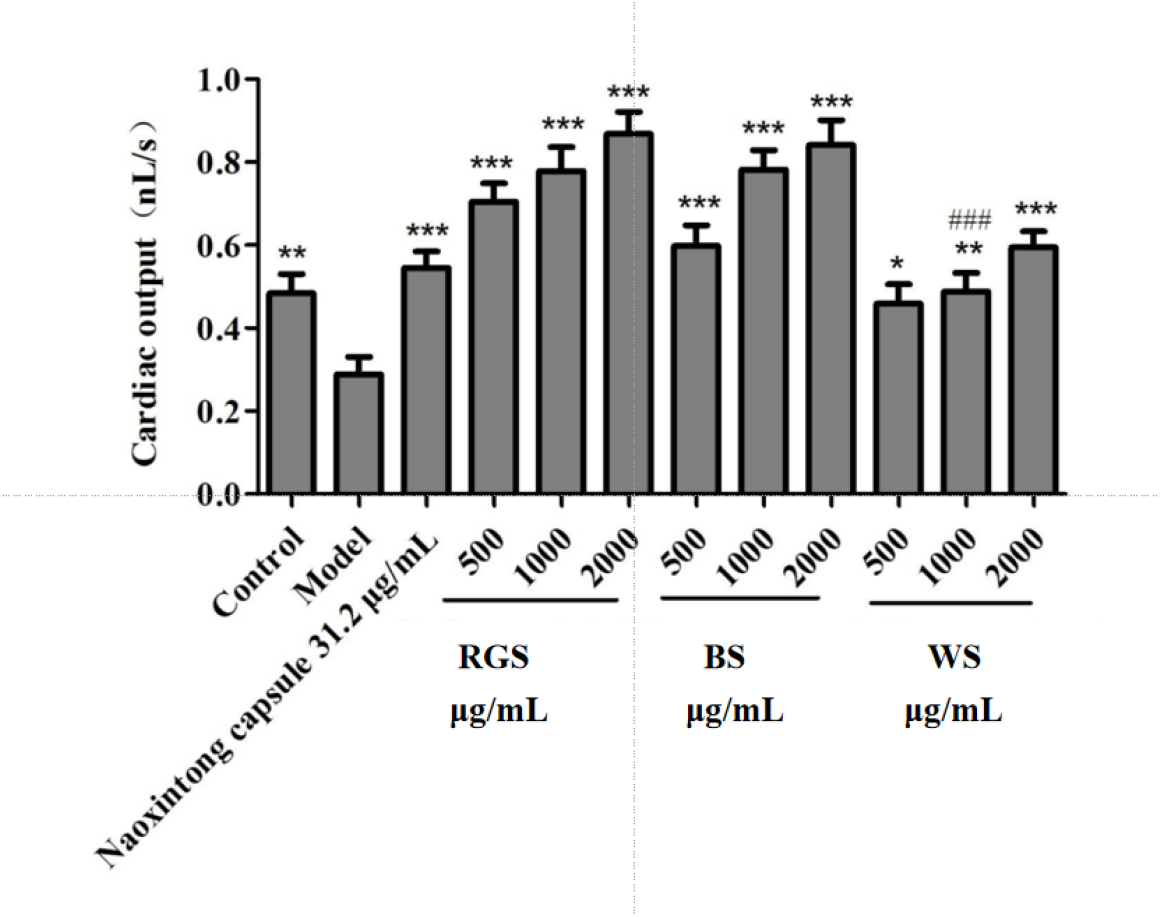
Cardiac output of zebrafish after sample treatment Compared with the model control group, *p < 0.05, **p < 0.01, ***p < 0.001 Compared with the 1000 µg/mL concentration group of ingamisin, ###p < 0.001

Under the experimental conditions, BS, RGS, and WS each demonstrated beneficial effects on qi deficiency and blood stasis, which were specifically reflected in the increased blood flow velocity and cardiac output. Notably, RGS and BS exhibited superior efficacy in improving these hemodynamic parameters compared to WS.

### 3.4 Evaluation of antioxidant effect in vivo

#### 3.4.1 Antioxidant effect (Yolk sac fluorescence intensity)

The zebrafish model is highly suitable for studying oxidative stress and antioxidant defense mechanisms due to its distinctive biological features, high genomic homology with humans, and antioxidant response pathways that resemble those in mammals. In zebrafish embryos, the yolk sac represents a critical organ whose fluorescence intensity is directly proportional to the level of reactive oxygen species (ROS) [14]. ROS are major byproducts of oxidative stress, and the antioxidant capacity of a test compound can be indirectly assessed by measuring yolk sac fluorescence intensity. As shown in Fig. 5 and 6, both BS and RGS exhibited significant antioxidant effects under the experimental conditions, whereas WS showed no detectable activity.

**Fig. 5.**
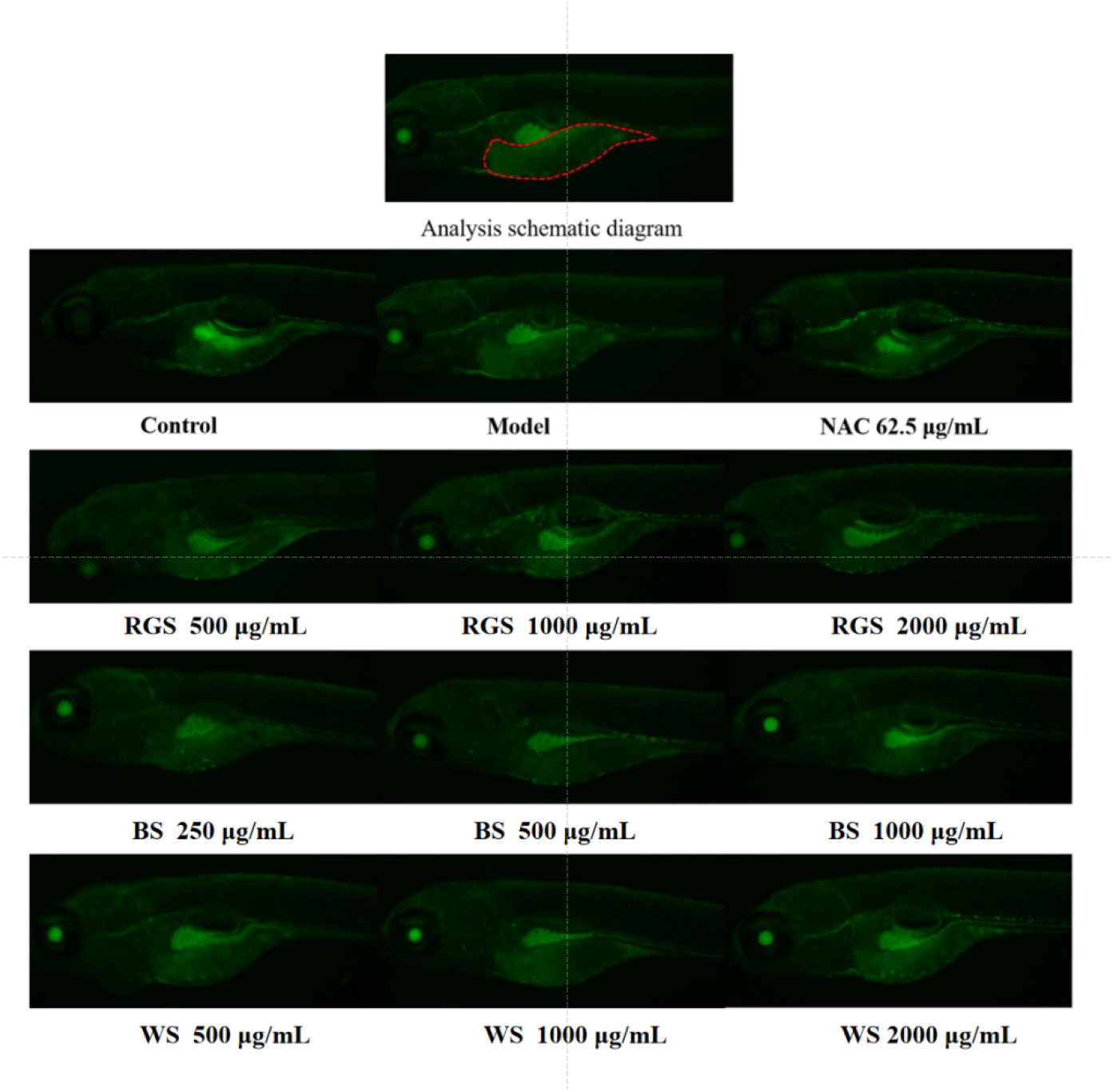
Typical fluorescence intensity of zebrafish yolk sac after sample processing Note: The red wireframe represents the analysis area

**Fig. 6.**
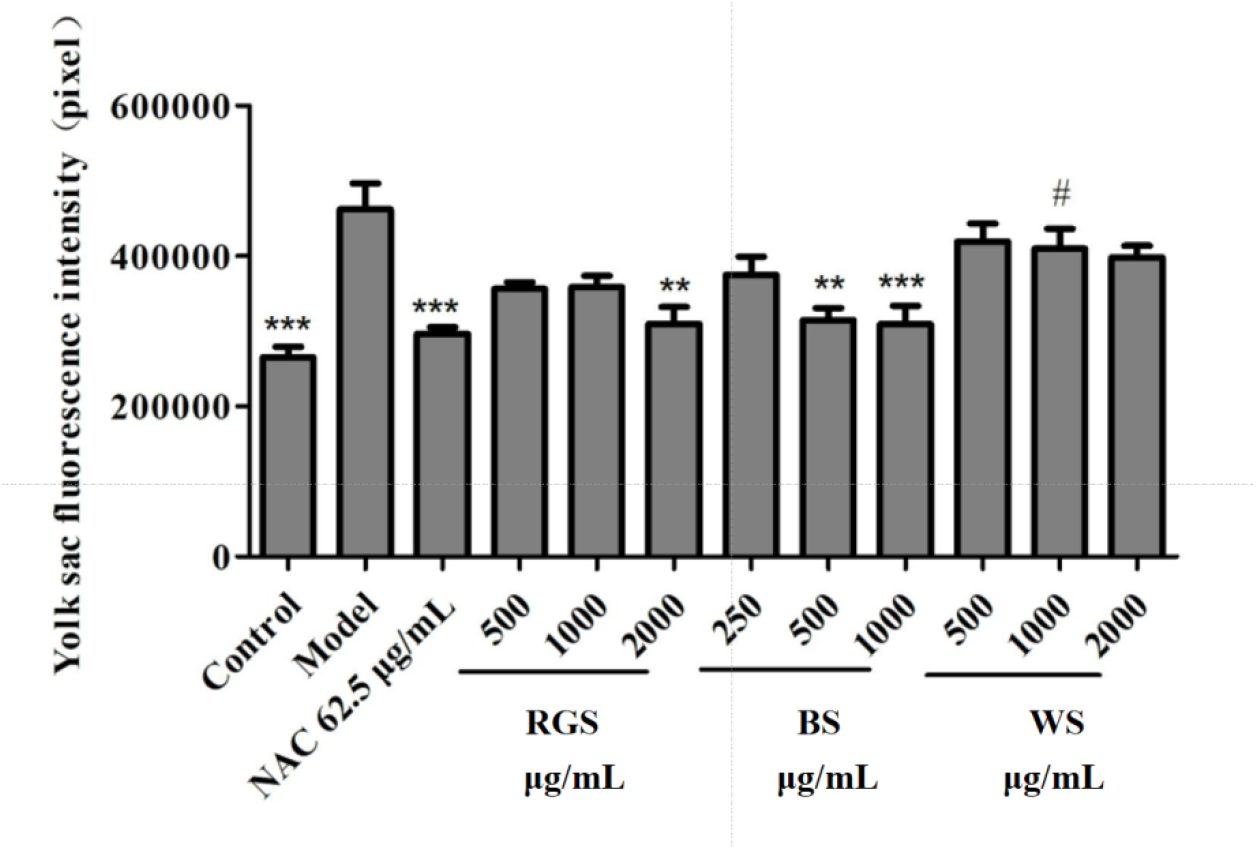
Fluorescence intensity of yolk sac after sample treatment Compared with the model control group, **p < 0.01, ***p < 0.001 Compared with the 1000 µg/mL group of inganose, #p < 0.05

#### 3.4.2 Antioxidant effect (SOD vitality)

The human antioxidant defense system comprises both enzymatic and non-enzymatic components. The enzymatic system primarily includes endogenous antioxidant enzymes such as superoxide dismutase (SOD) and catalase (CAT). As the primary defense against oxidative stress, SOD interrupts free radical chain reactions, thereby reducing free radical generation [15].

Using zebrafish embryos, we compared the in vivo antioxidant capacities of the three sugars. The results, presented in Fig. 7, indicate that at a concentration of 500 μg/mL, BS, RGS, and WS all significantly increased embryonic SOD activity compared to the control group (P < 0.05). Moreover, BS demonstrated a stronger effect on enhancing SOD activity than both RGS and WS.

**Fig. 7.**
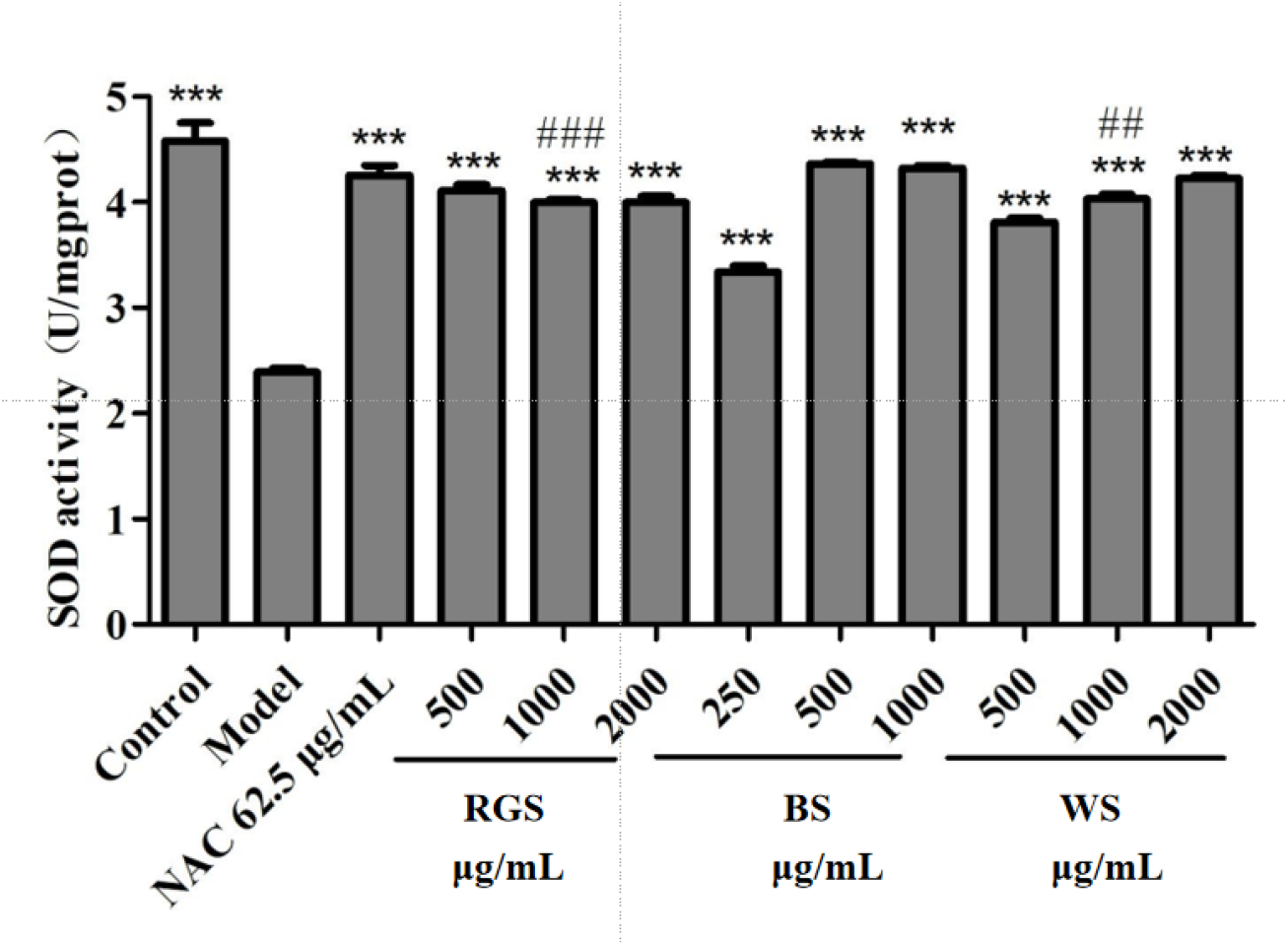
SOD activity after sample treatment Compared with the 1000 µg/mL group of inganose, ##p < 0.01, ###p < 0.001

### 3.5. Identification and quantification of nutrients in sugars

The efficacy of brown sugar may be attributed to its unique nutritional composition. Sugar processing methods vary considerably, leading to notable differences in the content of nutrients and trace elements [16], which in turn influence biological activity [17]. BS and RGS are non-centrifugal sugars that retain higher levels of sugarcane polyphenols, whereas WS is produced through centrifugal refining.

BS, a brown-red or yellow-brown sugar, is manufactured from sugarcane juice that is purified and boiled without centrifugation. Its short heating time helps preserve the nutritional and flavor profiles of sugarcane (in compliance with GB/T 35885-2018) [18]. RGS, a honey-brown or yellow-brown sugar, is also derived from sugarcane juice through extraction and purification. It retains some nutritional and flavor components but undergoes longer heating than BS (in compliance with GB/T 35884-2018) [19]. WS consists of crystalline sugar produced from sugarcane or sugar beet via juice extraction, purification, boiling, refining, and centrifugation (in compliance with GB 317-2018).

Quantitative analysis identified seven metallic elements, two non-metallic elements, and sugarcane polyphenols across all sugar samples (Fig. 8). BS contained all nine elements, whereas RGS and WS contained seven and five, respectively. BS showed the richest mineral composition, with calcium (Ca) and iron (Fe) levels more than three and five times those in RGS, respectively. Interestingly, RGS contained approximately four times more potassium (K) than BS.

**Fig. 8.**
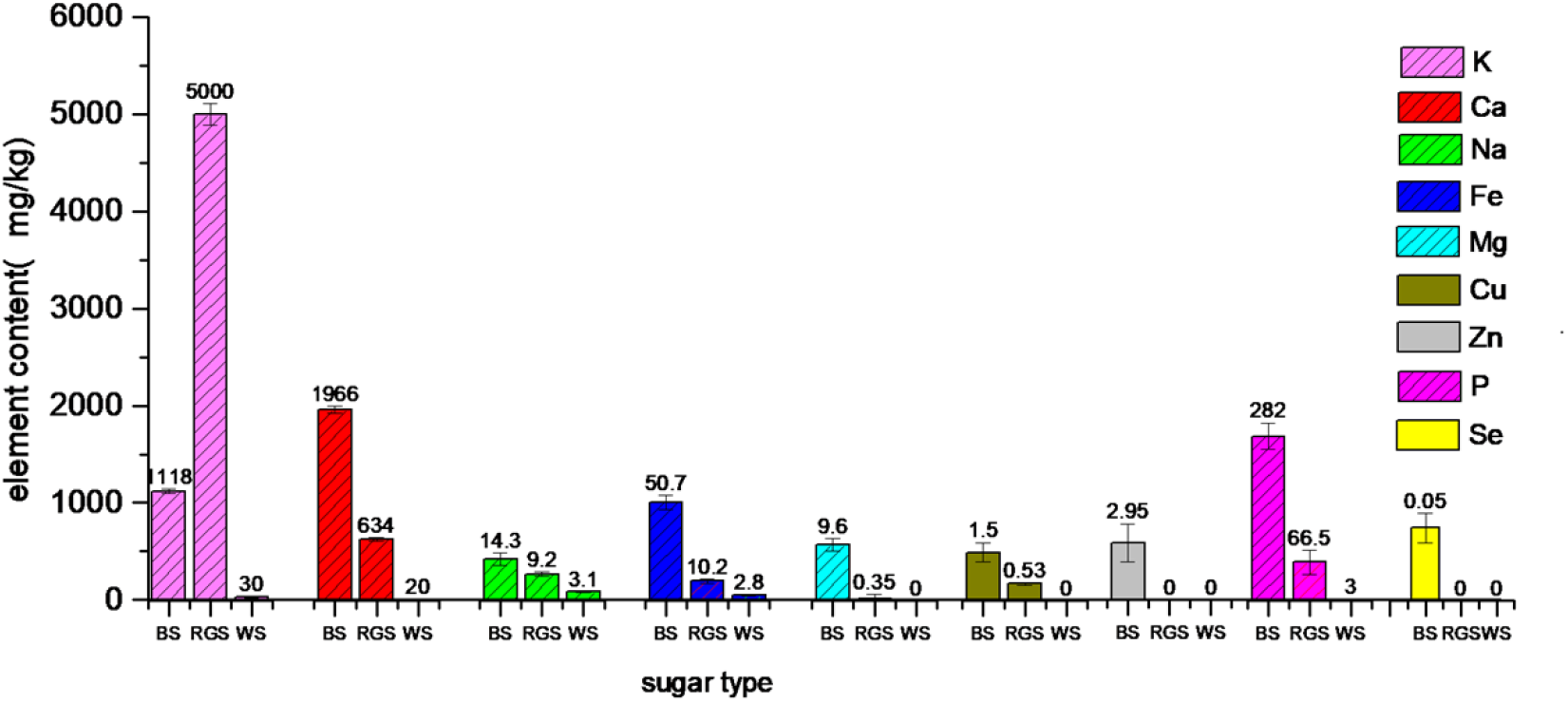
The nutritional elements content in three types of sugar

As shown in Fig. 9, BS had the highest sugarcane polyphenol content (4.35 g/kg), followed by RGS (1.43 g/kg); no polyphenols were detected in WS. These differences are attributable to variations in processing techniques [20]. Sugarcane polyphenols are natural antioxidants approved for use in food by the Chinese National Health Commission in 2022. Plant polyphenols constitute an important class of natural compounds with demonstrated functions such as antioxidant [21–24], anti-aging [25], anti-fatigue [26], hypoxia resistance [27], and potential therapeutic effects against age-related neurodegenerative diseases [28]. The antioxidant properties of sugarcane polyphenols likely contribute to the observed biological activities.

**Fig. 9.**
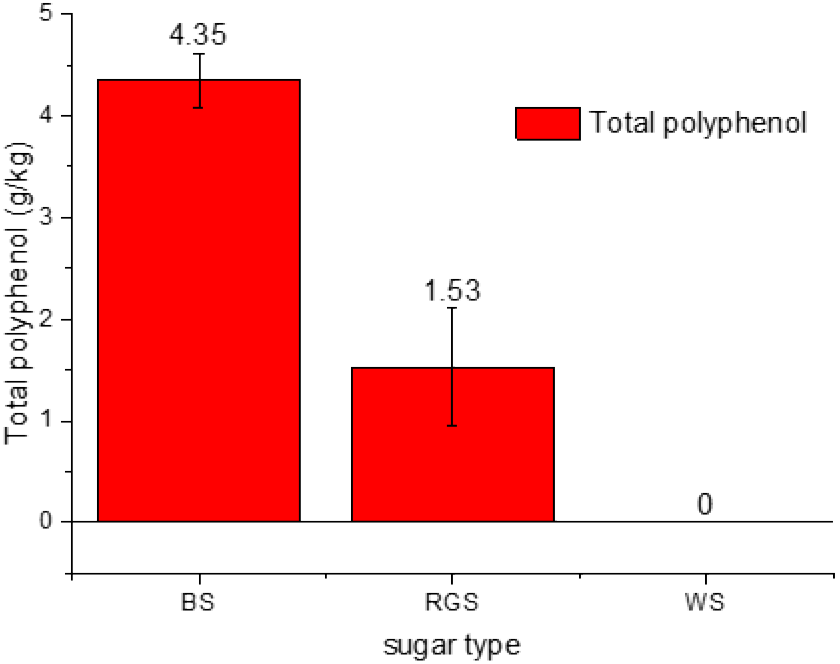
Content of total sugarcane polyphenol in different sugar

## 4 Discussion

BS contained significantly higher concentrations of nutritionally essential elements such as sodium, calcium, and iron compared to RGS and WS. These trace elements may underpin the enhanced functional performance of BS observed in zebrafish models.

In the context of physical fatigue alleviation, electrolytes (K^+^, Na^+^, Ca^2+^) help restore ionic balance, reducing the risk of post-exercise cramps and accelerating muscle recovery [29]. Trace elements such as copper, zinc, selenium, and vitamins in brown sugar may regulate metabolic processes in zebrafish, including glycogen synthesis and breakdown, thereby improving energy efficiency and reducing fatigue [30]. Iron, a key component of hemoglobin, enhances blood oxygen-carrying capacity and alleviates fatigue induced by hypoxia [31]. Furthermore, sugarcane polyphenols in BS and RGS may help eliminate exercise-induced ROS, reduce oxidative stress, and delay fatigue onset [32].

Modern lifestyles often lead to psychological stress and fatigue [33,34,35,36], which manifest not only as physical weakness but also as mental exhaustion, drowsiness, dyspnea, and generalized weakness [37]. Additional symptoms may include poor concentration, impaired memory, and low mood [38,39,40]. These findings provide a basis for developing brown sugar-based nutritional products aimed at mitigating fatigue.

Regarding the amelioration of qi deficiency and blood stasis, BS contained 50.7 mg/kg of iron—five times and eighteen times the levels in RGS and WS, respectively. Iron supplementation is known to improve iron-deficiency anemia [41], suggesting a mechanism for BS ‘ s efficacy. Elements such as selenium, magnesium, zinc, and vitamins in brown sugar may also promote blood circulation and improve symptoms related to qi and blood deficiency [42]. These components likely support hematopoiesis, enhance microcirculation, and resolve stasis. Minerals including calcium, sodium, and zinc contribute to vasodilation and improved vascular elasticity, further supporting circulatory health.

Brown sugar has a long history of use in traditional Chinese dietary therapy. Myocardial ischemia, characterized by reduced coronary blood flow and oxygen supply, disrupts cardiac energy metabolism [43]. Qi-tonifying and stasis-removing effects are integral to traditional Chinese management of coronary ischemia. With growing consumer interest in natural health products [44], future studies should explore the molecular mechanisms and synergistic effects of brown sugar with other blood-invigorating herbs to strengthen its scientific foundation.

The superior antioxidant activity of BS is likely due to its high polyphenol content, which scavenges free radicals, reduces ROS, and mitigates oxidative damage. In contrast, WS lacks these bioactive compounds, explaining its limited efficacy [45].

## 4. Conclusions

In vivo experiments demonstrated that brown sugar (BS) and raw granulated sugar (RGS) exhibited significantly stronger anti-fatigue effects and greater amelioration of qi deficiency and blood stasis compared to white sugar (WS) in zebrafish. Antioxidant assays confirmed that all three sugars significantly increased SOD activity in zebrafish embryos at 500 μg/mL. Overall, BS and RGS outperformed WS in antioxidant capacity, anti-fatigue benefits, and improvement of qi deficiency and blood stasis.

ICP analysis revealed nine nutritional elements (K, Ca, Na, Mg, Fe, Zn, Cu, P, Se) across the sugar types. BS contained all nine, with significantly higher P, Ca, Fe, and Mg than RGS. Selenium was detected only in BS. Polyphenol quantification showed that BS and RGS contained 4.35 g/kg and 1.53 g/kg of sugarcane polyphenols, respectively, while WS contained none. The superior nutrient and polyphenol profiles of BS and RGS are attributed to their less refined processing methods.

## Acknowledgements

This work was supported by Guangxi Science and Technology Major Project “Development and Industrialization Demonstration of Key Technologies for the Production and Application of High end Sucrose Products for Pharmaceutical Use” (**Project Number: Guike AA22117005**); Guangxi Key R&D Program Project “Development and Industrialization of Key Technologies for Quality Control of Medicinal Excipients Sucrose” (**Project Number: Guike AB22035032**); Chongzuo City Science Research and Technology Development Plan Project “Key Technology Development and Industrialization Demonstration of High end Injection Grade Medicinal Sucrose Product Production” (**Project Number: Chongke 2022QN1210**).

